# Comparisons among Vestibular Examinations and Symptoms of Vertigo in Sudden Sensorineural Hearing Loss Patients

**DOI:** 10.1101/479352

**Authors:** Kang Hyeon Lim, Yong Jun Jeong, Mun Soo Han, Yoon Chan Rah, Jaehyung Cha, June Choi

## Abstract

**Objective:** Vertigo in sudden sensorineural hearing loss (SSNHL) is hypothesized as an extension of the disease caused by the anatomical proximity of the cochlea and vestibule. The present study aimed to demonstrate the association of vestibular function test (VFT) results with SSNHL disease severity and prognosis.

**Subjects and methods:** This study assessed clinical records of 263 SSNHL patients admitted to our hospital, between January 2010 and October 2017. Steroid treatment comprised high-dose intravenous dexamethasone (16 mg/d) or oral methylprednisolone (64 mg/d) for 4 days and tapered oral methylprednisolone for 8 days after discharge. Caloric tests were performed in all patients, and cervical vestibular-evoked myogenic potential (c-VEMP) and ocular VEMP (o-VEMP) tests were performed in 209 and 144 patients, respectively.

**Results:** Ninety six patients had vertigo, and caloric abnormalities were observed in 119 patients. Initial PTA in patients with vertigo were worse than in those without vertigo (63.0 dB vs 72.7 dB, *P*=.002). Initial PTA in patients with abnormal o-VEMP was worse than in those with normal o-VEMP (61.4 dB vs 73.0 dB, *P*=.004). PTA improvement after steroid treatment in patients with vertigo was lower than in those without vertigo (25.0 dB vs 20.9 dB, *P*=.028). PTA improvement after treatment in patients with abnormal caloric results was lower than in those with normal caloric results (26.0 dB vs 18.4 dB, *P*=.013).

**Conclusions:** The functions of vestibular organs, particularly the utricle and lateral semicircular canal, are associated with disease severity and hearing outcome in SSNHL patients.

## Introduction

Sudden sensorineural hearing loss (SSNHL) is clinically defined as sensorineural hearing loss of over 30 dB in more than 3 contiguous audiometric frequencies, occurring within 3 d [1]. Tinnitus occurs in approximately 80% of patients and vertigo in approximately 30% to 40% [2,3]. Vertigo appears more frequently in association with profound hearing loss than with mild or moderate deafness, and hearing recovery is worse in patients with vertigo than it is in those without vertigo [3–5].

Vestibular involvement in SSNHL is hypothesized as an extension of the disease caused by the anatomical proximity and correlation of the cochlea and vestibule [6]. Recently, vestibular diagnostic methods have been included in the general evaluation of SSNHL patients, such as the caloric test, cervical vestibular-evoked myogenic potential (c-VEMP), and ocular VEMP (o-VEMP) [7]. The caloric test is a method for investigating lateral semicircular canal (LSCC) function and superior vestibular integrity, the c-VEMP can be used for assessing saccular function and the inferior vestibular pathway, and the o-VEMP can be used for evaluating utricular function and the superior vestibular pathway [8–11].

Regarding the findings of vestibular function tests (VFTs) in SSNHL with vertigo, 40% of patients have previously been reported to show reduced caloric response in the affected ear.[1,4,5,12] Some reports have indicated that abnormal caloric test results in SSNHL patients may be of negative prognostic value [1,13], whereas other studies have shown that the results of caloric testing may not be predictive of SSNHL [4,5,14]. Furthermore, several recent studies have included the VEMP test in the evaluation of patients with SSNHL, for whom abnormal results of vestibular examinations were associated with profound hearing loss [6,15].

The present study aimed to determine the association of VFT results with the initial disease severity and prognosis of hearing outcomes in SSNHL patients.

## Materials and Methods

### Study Patients

We retrospectively reviewed the clinical records of 263 patients with SSNHL who had been admitted to the Department of Otorhinolaryngology-Head and Neck Surgery of our hospital between January 2010 and October 2017. All patients had sudden-onset idiopathic unilateral sensorineural hearing loss, which was defined as a 30-dB loss over 3 contiguous frequencies occurring within 3 d. Patients who presented with chronic otitis media or inner ear abnormalities on magnetic resonance imaging, and a history of surgery in the affected ear were excluded. Initial evaluations, including physical examination, pure tone audiometry, and VFTs, were performed within 14 d after onset and treatment was started immediately. Steroid treatment comprised high-dose intravenous dexamethasone (16 mg/d) or oral methylprednisolone (64 mg/d) for 4 d during hospitalization and tapered oral methylprednisolone for 8 d after discharge. This study was approved from Institutional review board approval of our hospital (IRB No: 2018AS0093).

### Audiometry and Hearing Outcome

Repeated pure tone audiometry was conducted every other day for each admitted patient. Pure tone averages (PTAs) were calculated by averaging the pure tone levels at 500 Hz, 1000 Hz, 2000 Hz, and 3000 Hz. Initial hearing loss was classified in 5 degrees by air conduction: mild (PTA 20–39 dB), moderate (PTA 40–59 dB), severe (PTA 60–79 dB), profound (PTA 80–99 dB), and deaf (PTA ≥100 dB). One month after the steroid treatment, patients underwent follow-up pure tone audiometry, and the hearing outcome was assessed according to Siegel’s criteria: 1) Complete recovery to a final hearing level better than 25 dB regardless of the initial hearing, 2) Partial recovery by more than 15 dB of gain or a final hearing level between 25 and 45 dB, 3) Slight improvement by more than 15 dB of gain and a final hearing level worse than 45 dB, 4) No improvement, which was classified as less than 15 dB of gain and a final hearing level worse than 75 dB [16].

### Vestibular Function Test

All patients underwent a bithermal caloric test, and 209 and 144 patients had c-VEMPs and o-VEMPs, respectively (Fig 1). Caloric tests were performed by irrigating the external auditory canal with water at 30°C and 44°C. The maximum slow phase velocity of nystagmus induced by caloric stimulation of the right or left side was recorded. Canal paresis (CP) was calculated as a percentage by using Jonkee’s formula: CP = 100{(right side maximum slow-phase velocity – left side maximum slow-phase velocity) / (right side maximum slow-phase velocity + left side maximum slow-phase velocity)} CP of more than 20% was regarded as abnormal [17].

**Fig 1.**
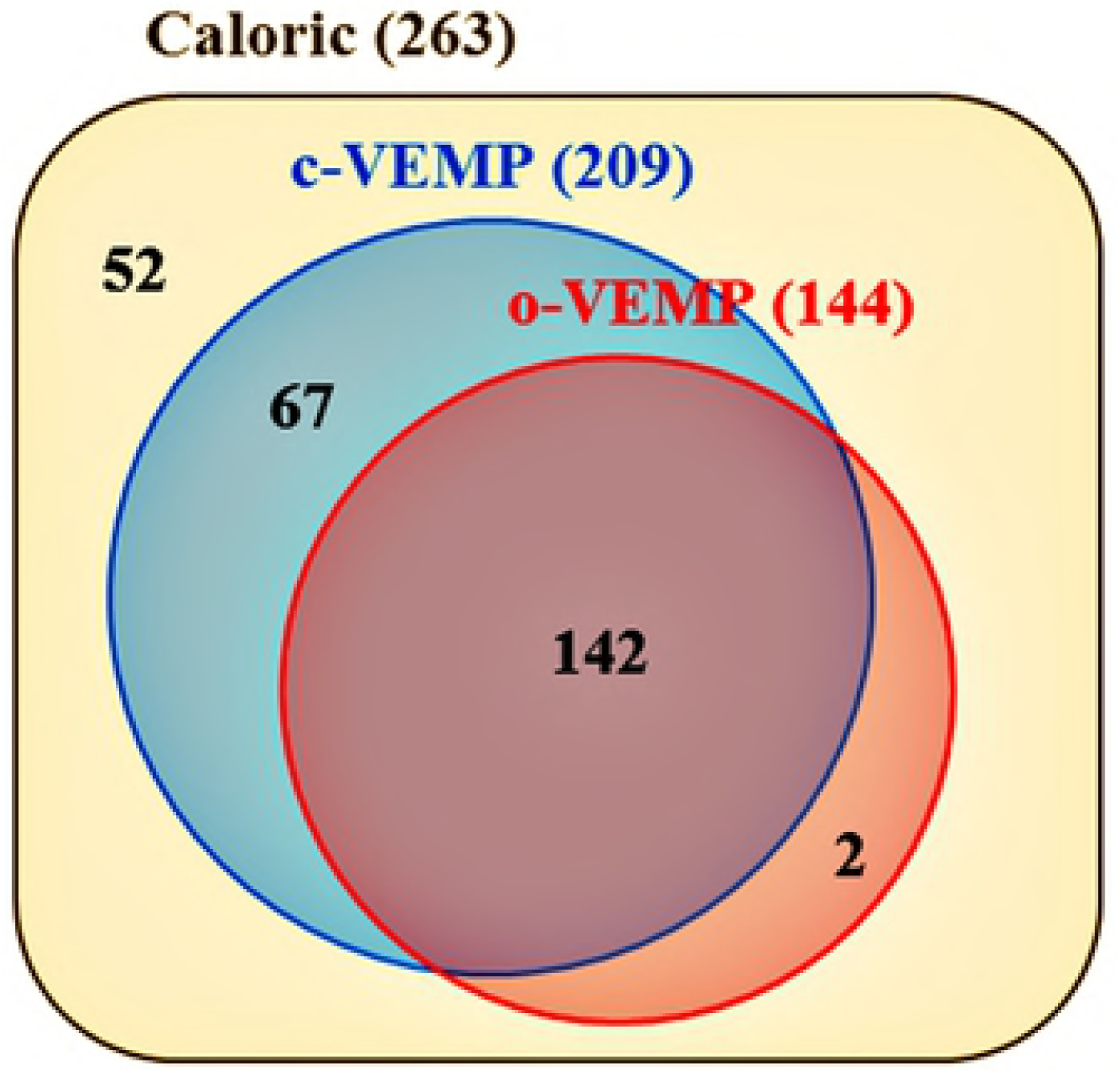
Analysis of vestibular function test. Bithermal caloric tests were performed in 263 patients, and cervical vestibular-evoked myogenic potential (c-VEMP) and ocular vestibular-evoked myogenic potential tests were performed in 209 and 144 patients, respectively.

During c-VEMP testing, electromyographic (EMG) activity was recorded from a surface electrode placed on the upper half of each sternocleidomastoid muscle (SCM), with a reference electrode on the side of the upper sternum and a ground electrode on the chin. The patients were asked to raise their heads in the supine position to make contract with the SCM. Acoustic stimuli were delivered through headphones. The first positive wave form deflection was marked as p13 and the first negative deflection was marked as n23. The latencies and amplitudes of these waveforms (p13-n23) were measured on both sides. For the evaluation of amplitude, the interaural amplitude difference ratio (IAD) was calculated as a percentage as follows: 100{(Au – Aa) / (Aa + Au)}, where Au is the p13–n23 amplitude on the unaffected side and Aa is the p13–n23 amplitude on the affected side.[18] An IAD in c-VEMP amplitude of more than 20% or an absence of amplitude on either side was regarded as an abnormal c-VEMP result.

For o-VEMP, patients adopted a sitting position with surface EMG electrodes placed 1 cm below (active) and 3 cm below (indifferent) the center of each lower eyelid. The ground electrode was placed on the chin. The patients looked up, approximately 30° above straight ahead, and maintained their focus on a small dot approximately 1 m from their eyes. The first negative peak (n10) latency, the subsequent positive peak (p16) latency, and the amplitude between n10 and p16 were recorded from the eye contralateral to the stimulation. For the evaluation of amplitude, the IAD for n10–p16 amplitude was calculated as a percentage, as follows: 100{(Au – Aa) / (Aa + Au)}, where Au is the n10–p16 amplitude on the unaffected side and Aa is the n10–p16 amplitude on the affected side.[19] An IAD in o-VEMP amplitude of more than 20% or an absence of amplitude on either side was regarded as abnormal.

### Statistical Analysis

The Statistical Package for the Social Sciences Version 21 (IBM Corporation, NY, USA) was used for statistical analysis. The chi-square test and Kruskal–Wallis test were used to compare clinical characteristics among the 5 groups according to the pretreatment hearing levels. The Mentel–Haenszel trend test was used for assessing the trend of categorical variables, and the Mann–Whitney test was used for comparing continuous variables. A difference of *P*<.05 was regarded as significant.

## Results

No significant differences were observed in the ratios of sex, diabetes mellitus, hypertension, and dyslipidemia among the 5 groups classified according to the pretreatment hearing levels, and the mean ages of the groups did not differ significantly. Of the 263 patients with SSNHL, including 75 men and 188 women, symptoms of vertigo were present in 96 patients (36.5%), and abnormal caloric responses were observed in 119 patients (45.2%). Diabetes mellitus, hypertension, and dyslipidemia were present in 53 (20.2%), 67 (20.2%), and 13 (4.9%) patients, respectively (Table 1). The mean initial PTA was worse in the patients with vertigo (72.7 dB) than in the patients without vertigo (63.0 dB) (*P*=.002), and the abnormal o-VEMP group had a worse mean initial PTA (73.0 dB) than the normal o-VEMP group (61.4 dB) (*P*=.004). However, no significant difference was observed in initial hearing loss severity between the normal and abnormal caloric response groups (*P*=.744), or between the normal and abnormal c-VEMP groups (*P*=.059) (Fig 2a).

**Table 1.** Clinical characteristics of 5 groups according to the pretreatment hearing levels (%).

| Variable | Total<br>(n=263) | Mild<br>(n=52) | Moderate<br>(n=57) | Severe<br>(n=55) | Profound<br>(n=46) | Deaf<br>(n=53) | *P*-value |
| --- | --- | --- | --- | --- | --- | --- | --- |
| Sex |  |  |  |  |  |  | .575<sup>†</sup> |
| Male | 75 (28.5%) | 11 (21.2%) | 17 (29.8%) | 14 (25.5%) | 16 (34.8%) | 17 (32.1%) |  |
| Female | 188 (71.5%) | 41 (78.8%) | 40 (70.2%) | 41 (74.5%) | 30 (65.2%) | 36 (67.9%) |  |
| Age, mean (SD), y | 48.9 (12.8) | 16.4 (12.5) | 47.8 (13.4) | 50.1 (11.8) | 48.6 (14.2) | 21.6 (12.4) | .653<sup>‡</sup> |
| Diabetes mellitus |  |  |  |  |  |  | .241<sup>†</sup> |
| Absent | 210 (79.8%) | 45 (86.5%) | 45 (78.9%) | 44 (80.0%) | 39 (84.8%) | 37 (69.8%) |  |
| Present | 53 (20.2%) | 7 (13.5%) | 12 (21.1%) | 11 (20.0%) | 7 (15.2%) | 16 (30.2%) |  |
| Hypertension |  |  |  |  |  |  | .193<sup>†</sup> |
| Absent | 196 (74.5%) | 40 (76.9%) | 42 (73.7%) | 44 (80.0%) | 37 (80.4%) | 33 (62.3%) |  |
| Present | 67 (25.5%) | 12 (23.1%) | 15 (26.3%) | 11 (20.0%) | 9 (19.6%) | 20 (37.7%) |  |
| Dyslipidemia |  |  |  |  |  |  | .296<sup>†</sup> |
| Absent | 250 (95.1%) | 51 (98.1%) | 55 (96.5%) | 51 (92.7%) | 45 (97.8%) | 48 (90.6%) |  |
| Present | 13 (4.9%) | 1 (1.9%) | 2 (3.5%) | 4 (7.3%) | 1 (2.2%) | 5 (9.4%) |  |
| Vertigo |  |  |  |  |  |  | .001<sup>†</sup> |
| Absent | 167 (63.5%) | 37 (71.2%) | 37 (64.9%) | 43 (78.2%) | 28 (60.9%) | 22 (41.5%) |  |
| Present | 96 (36.5%) | 15 (28.8%) | 20 (35.1%) | 12 (21.8%) | 18 (39.1%) | 31 (58.5%) |  |
| Caloric response |  |  |  |  |  |  | .190<sup>†</sup> |
| Normal | 144 (54.8%) | 27 (51.9%) | 30 (52.6%) | 34 (61.8%) | 30 (65.2%) | 23 (43.4%) |  |
| Abnormal | 119 (45.2%) | 25 (48.1%) | 27 (47.4%) | 21 (38.2%) | 16 (34.8%) | 30 (56.65) |  |
| Treatment starting<br>from onset, mean<br>(SD), d | 4.11 (4.33) | 4.5 (4.3) | 4.9 (4.6) | 5.1 (5.4) | 2.9 (2.1) | 2.6 (3.5) | .030<sup>‡</sup> |

<sup>†</sup>Chi-square test was used.
<sup>‡</sup>Kruskal–Wallis test was used.
SD = standard deviation

**Fig 2.**
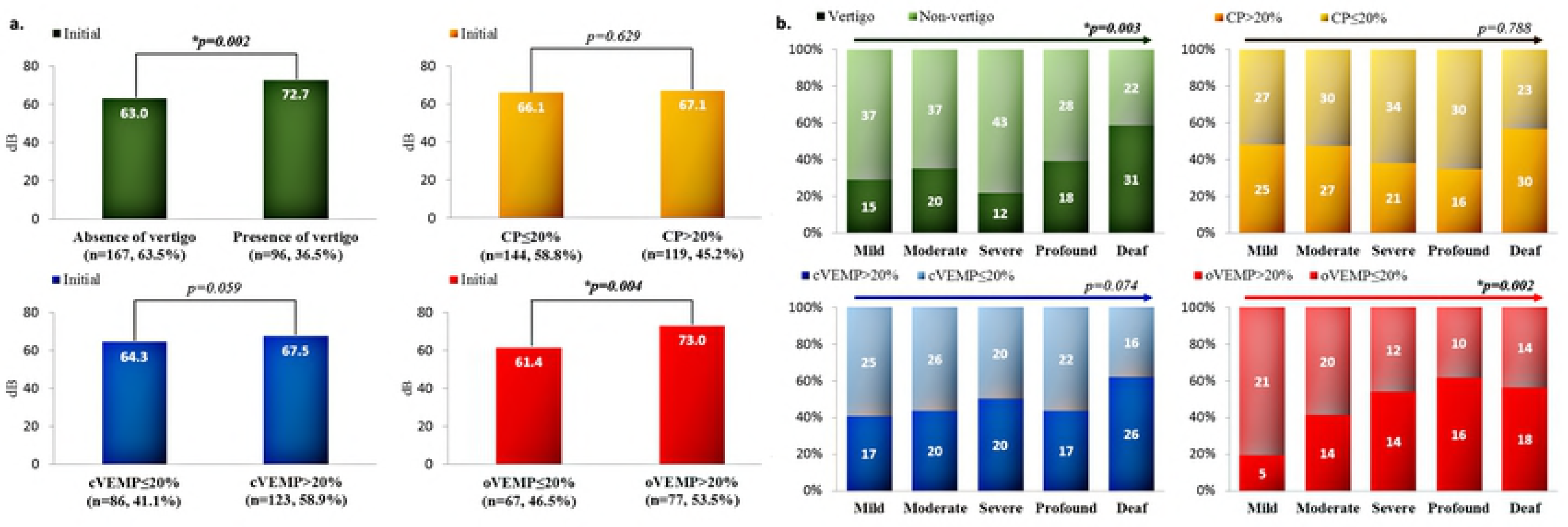
Relationship of initial hearing and vestibular function. Mean pure tone averages before treatment were compared between patients with and without vertigo, and between patients with normal and abnormal vestibular function test results. The Mann–Whitney test was used to determine statistical significance in the comparison of the symptom of vertigo (*P*=.002) and in the ocular vestibular-evoked myogenic potential (o-VEMP) (*P*=.004) (a). Trends in the proportion of vertigo and abnormal vestibular function test results among the 5 groups, classified according to the initial pure tone averages. The Mentel–Haenszel trend test was used to determine a significant increase in the proportion of vertigo (*P*=.003) and o-VEMP abnormality (*P*=.002) among the disease severities (b).

Furthermore, the 5 groups were classified according to the initial PTA and analyzed in terms of vertigo, canal paresis, c-VEMP, and o-VEMP. As initial hearing loss severity worsened from mild to deaf, the ratio of patients with vertigo and abnormal o-VEMP significantly increased (*P*=.003, *P*=.002). No significant trend was observed in the ratio of abnormal caloric response or c-VEMP abnormality among the groups (*P*=.788, *P*=.074) (Fig 2b).

The mean improvement in PTA (*Δ_mean_*) after steroid treatment was significantly lower in the patients with vertigo than in the patients without vertigo (*P*=.028), and the patients with abnormal caloric response exhibited a lower improvement in mean PTA than those with normal caloric response (*P*=.013). However, no significant difference was observed in hearing improvement between the normal and abnormal c-VEMP or o-VEMP groups (*P*=.118, *P*=.060) (Fig 3a).

**Fig 3.**
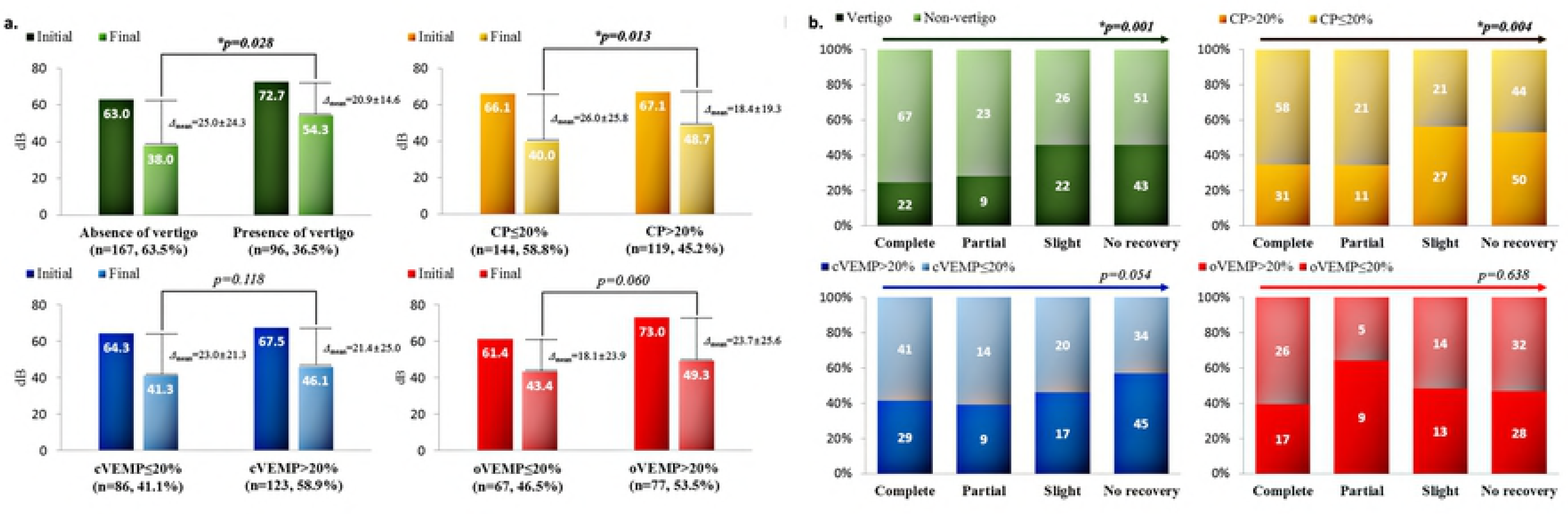
Relationship of hearing improvement and vestibular function. The mean difference of initial and final pure tone averages (*Δ_mean_*) was compared between patients with and without vertigo, and between patients with normal and abnormal vestibular function test results. The Mann–Whitney test was used to determine statistical significance in the comparison of the symptom of vertigo (*P*=.028) and in caloric test results (*P*=.013) (a). Trends in the proportion of vertigo and abnormal vestibular function test results among the 4 groups classified according to Siegel’s criteria. The Mentel–Haenszel trend test was used to determine a significant increase in the proportion of vertigo (P=.001) and caloric test abnormality (*P*=.004) among the hearing outcomes (b).

Analysis of the groups classified according to Siegel’s criteria revealed that as the treatment response of the groups worsened form complete to no recovery, the ratio of patients with vertigo and abnormal caloric response significantly increased (*P*=.001, *P*=.004) [16]. However, no significant trend was observed in the ratio of abnormal c-VEMP or o-VEMP among the groups (*P*=.054, *P*=.638) (Fig 3b).

## Discussion

Since SSNHL with vestibular involvement was first reported in 1949, the symptom of vertigo has been found to be associated with severe hearing loss and poor prognosis [3–5,20]. Although vertigo in SSNHL is hypothesized to be an extension of the disease from the cochlea to the vestibular organs because of their anatomical proximity, the relationship and pathogenesis of cochlear dysfunction and vestibular involvement remain controversial [6].

In the present study, impaired vestibular functions, as well as the symptom of vertigo, were found to be associated with hearing loss severity and prognosis. Utricular dysfunction, monitored using the o-VEMP test, demonstrated a relationship with severe hearing loss and dysfunction of the LSCC, and monitored using the caloric test, demonstrated a relationship with poor prognosis. In a meta-analysis evaluating vestibular function in SSNHL, vestibular organs were found to be involved in SSNHL regardless of the presence of vertigo, indicating that vertigo might not be an independent and determining factor. Vestibulocochlear lesion patterns showed that the utricle, followed by the LSCC, was the most susceptible to damage in SSNHL, and abnormal caloric response was a negative prognostic marker [7].

Thrombosis and vasospasm have been introduced as possible vascular mechanisms to explain isolated labyrinthine infarction [21]. Sudden vertigo and hearing loss are common accompanying symptoms in anterior inferior cerebellar artery (AICA) infarctions [22,23]. Decreased blood flow in the vertebrobasilar system caused by occlusive disease may selectively damage the inner ear because it has high energy requirements and lacks collaterals [24]. Spotty degeneration in the end organs can be produced by interruption of blood flow [25].

The internal auditory artery (IAA) typically originates from the AICA and supplies blood to the inner ear [26]. Within the internal auditory canal, the IAA divides into two main branches: the common cochlear artery and the anterior vestibular artery. The common cochlear artery further bifurcates into the main cochlear artery and the vestibulocochlear artery, with the latter forming the posterior vestibular artery and the cochlear ramus. The main cochlear artery and cochelar ramus respectively irrigate apical three fourths and basal one fourth of the cochlea. The utricle, superior part of the saccule, and ampullae of the superior and lateral SCC are supplied with blood by the anterior vestibular artery, while the inferior part of the saccule and the ampulla of the posterior SCC are supplied by the posterior vestibular artery (Fig 4) [27].

**Fig 4.**
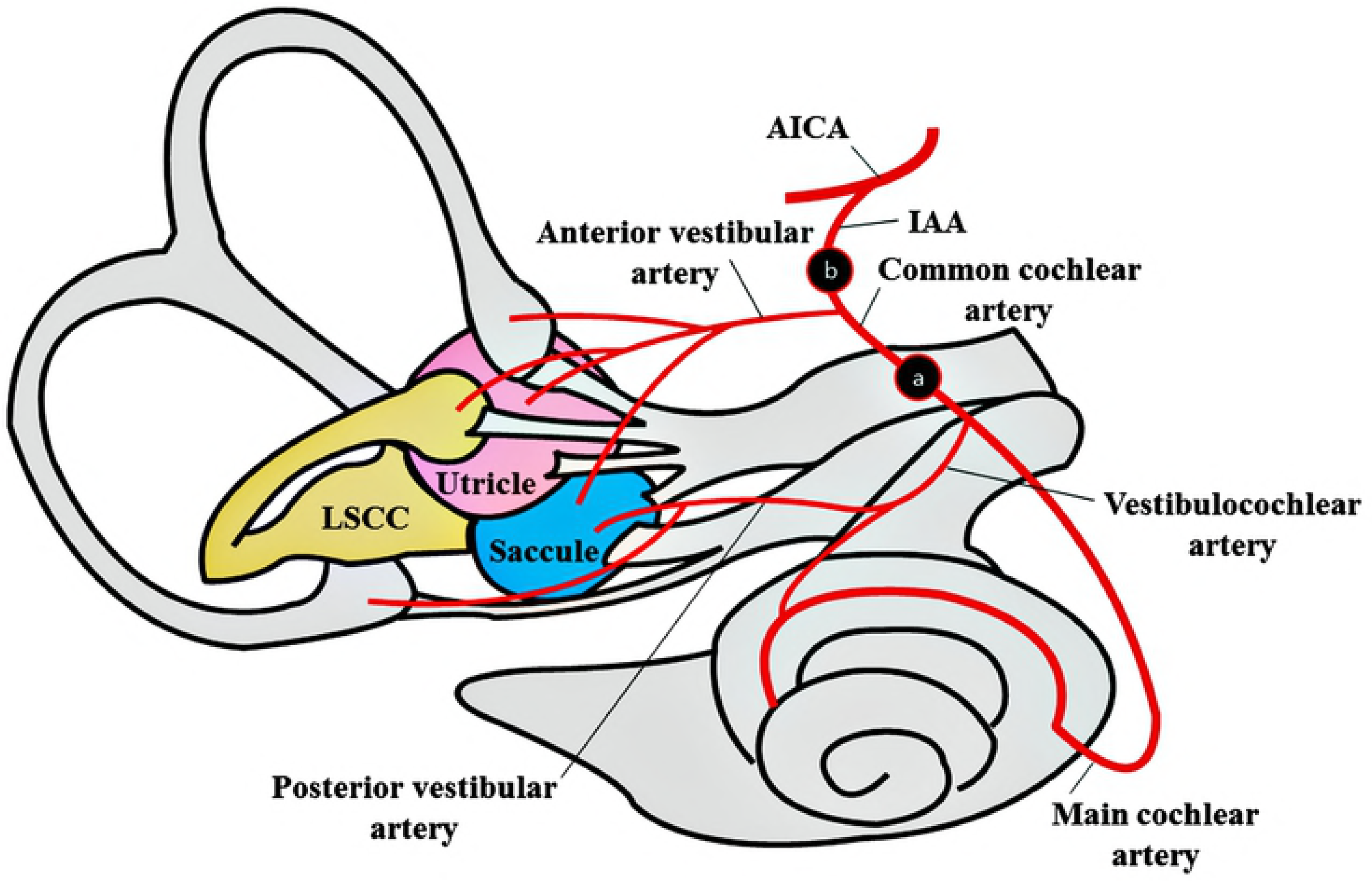
Vascular hypothesis in sudden sensorineural hearing loss. Because the cochlea and saccule are mainly supplied with blood from the branches of the common cochlear artery, infarction therein may lead to cochlear and saccular dysfunction. The saccule may be more resistant to ischemic damage because of its intraosseous collateral branches (a). Infarction in the internal auditory artery may lead to hypoperfusion of the anterior vestibular artery and common cochlear artery, which in turn leads to dysfunction of the cochlea, utricle, and lateral semicircular canal. The utricle and lateral semicircular canal are considered to be more vulnerable to ischemic damage than the saccule (b).

Whereas an abnormal o-VEMP was related to severe initial hearing loss, and an abnormal caloric response was related to a poor hearing outcome in SSNHL patients, c-VEMP results were not significantly related to hearing loss severity or prognosis in this study. Becuase the c-VEMP reflects saccular function and perfusion to the saccule is mainly supplied by the posterior vestibular arteries, this could be explained by differential effects of ischemia on the anterior and posterior vestibular arteries. Given that intraosseous collaterals of the vestibulocochlear artery and posterior vestibular artery are more abundant than those of the anterior vestibular artery, the saccule may be more resistant to ischemic damage and saccular function may be a less accurate reflection of the degree of ischemia of the cochlea than the utricle and LSCC.[28] Furthermore, although the cochlea and saccule are mainly supplied with blood from the branches of the common cochlear artery, saccular function may not worsen as much as cochlear function in common cochlear artery infarction (Fig 4).

The function of the utricle or LSCC could be impaired when perfusion to the anterior vestibular artery becomes insufficient. Dysfunction of the utricle or LSCC in SSNHL patients could imply IAA infarction because perfusions of the cochlear artery and anterior vestibular artery are impaired simultaneously. Therefore, SSNHL patients with abnormal o-VEMP or caloric results may have a broader territory of infarction, including the anterior vestibular artery, than those with normal o-VEMP and caloric results. In this study, o-VEMP and caloric abnormality were associated with disease severity and poor prognosis, respectively, which may be due to the broad extent of the disease involving the utricle and LSCC. The reason o-VEMP results were associated with an immediate worsening of severity and caloric results were associated with delayed prognosis may be that artery supplying the utricle is more proximally branched out from the anterior vestibular artery than the artery supplying the LSCC. Utricular function reflecting the effect of IAA occlusion more early than the function of the LSCC is a possible explanation.

The present study has some limitations. Selection bias and information bias could have occurred because patients were analyzed retrospectively. Although the results could have been affected by them, the type of hearing loss, the degree of vertigo, and other accompanying symptoms were not analyzed in this study. Finally, the patient population was not identical in terms of caloric, c-VEMP, and o-VEMP tests.

## Conclusions

The functions of vestibular organs are associated with disease severity and have predictive value in hearing outcomes in SSNHL patients with or without vertigo. The wider the extent of disease is, particularly when it involves the LSCC, the poorer the hearing outcome is. The pattern of vestibular organ dysfunction correlates with the distribution of the branches of the IAA. Thus, the vascular hypothesis of the inner ear should be further studied to explain the pathogenesis of SSHNL and vestibular involvement.

## Competing interests

The authors have declared that no competing interests exist.

## Author Contributions

Conceptualization: Kang Hyeon Lim, Yong Jun Jeong, Mun Soo Han, June Choi

Data curation: Kang Hyeon Lim, Yoon Chan Rah, Jaehyung Cha, June Choi

Formal analysis: Kang Hyeon Lim, Mun Soo Han, Jaehyung Cha, June Choi

Investigation: Kang Hyeon Lim, Yong Jun Jeong, Mun Soo Han, June Choi

Methodology: Kang Hyeon Lim, June Choi

Validation: Kang Hyeon Lim, Yoon Chan Rah, Jaehyung Cha, June Choi

Writing ± original draft: Kang Hyeon Lim, June Choi

Writing ± review & editing: Kang Hyeon Lim, June Choi

